# Pointing in depth is shaped by a natural grasping distance prior

**DOI:** 10.1101/2020.12.11.421206

**Authors:** Michael Wiesing, Tatiana Kartashova, Eckart Zimmermann

**Author notes:** Corresponding author: Michael Wiesing.

## Abstract

Vision in depth is distorted. A similar distortion can be observed for pointing to visual targets in depth. It has been suggested that pointing errors in depth reflect the visual distortion. However, much research has suggested that in case visual information is not rich enough, the sensorimotor system involves prior knowledge to optimally plan movement trajectories. Here, we show that pointing in depth is guided by a prior that biases movements toward the natural grasping distance at which object manipulation is usually performed. To dissociate whether pointing is guided by distorted vision only or whether it takes into account a natural grasping distance prior, we adapted pointing movements. Participants received visual feedback about the success of their pointing once the movement was finished. We distorted the feedback to signal either that pointing was not far enough or in separate sessions that pointing was too far. Participants adapted to this artificial error by either extending or shortening their pointing movements. The generalization of pointing adaptation revealed a bias in movement planning that is inconsistent with pointing being guided only by distorted vision but with the involvement of knowledge about the natural grasping distance. Adaptation was strongest for pointing movements to a middle position that corresponds to the natural grassing distance and it was weakest for movements leading away from it. It has been demonstrated that pointing adaptation in depth changes visual perception (Volcic et al., 2013). We also wondered how effects of pointing adaptation on visual space would generalize in depth.

## Introduction

How do we determine the locations of objects in three-dimensional space? The primary visual cues to depth – in the absence of motion parallax – are binocular cues. Although monocular cues can in principle guide movements, reaching performance in one-eyed vision drops down dramatically, suggesting the dominance of binocular cues in natural depth vision (Servos et al., 1992; Marotta er al., 1995). However, depth information provided by binocular disparity can only be interpreted ambiguously such that a small object that is close produces the same cues as a large object that is far away. When compensating this ambiguity through ocular convergence, the brain generates systematic errors in depth perception: Objects near to an observer are overestimated and objects far away are underestimated (Johnston, 1990; Norman et al., 1996).

Pointing in depth is similarly distorted: We overshoot close objects and undershoot far objects (Bradshaw & Hibbard, 2003). It seems natural to assume that this pointing behavior is the consequence of the visual distortion. However, for two reasons it would be surprising if the brain left the sensory ambiguity uncompensated. First, many experimental findings have corroborated the proposal of a distinct neural processing of vision-for-perception and vision-for-action and have demonstrated that arm movements remain accurate even if vision is incorrect (for a review, see Goodale, 2014). Second, much research highlights that sensorimotor performance is optimized by weighing prior knowledge with sensory information in a near-optimal way (e.g. Körding and Wolpert 2004; Tassinari et al. 2006; Vilares et al. 2012). In the absence of precise visual information, consulting prior knowledge supports the choice of a movement that maximizes utility (Körding and Wolpert, 2006). For sensorimotor decisions, utility is a weighed comprise between the likely visual target position and the cost of the movement. In case visual information is weak, movements will be guided stronger by priors about movements costs that have been built up over the lifetime (Berniker et al. 2010). For arm movements, the “natural grasping distance” describes the location of the hand where object manipulation is performed most efficiently (Volcic, 2013). Here, we asked if humans compensate the 3D ambiguity in visual depth perception by employing a “natural grasping distance” prior. To this end, we implemented pointing adaptation experiments that could dissociate whether the generalization of adaptation follows visual distortions or if it is informed by a “natural grasping distance” prior. Observers pointed to visual targets in a virtual environment without seeing their hands and received distorted feedback about the success of the movement when they reached the target.

We also measured how pointing adaptation might affect visual space. Volcic et al. (2013) have provided evidence that pointing adaptation can shift the point where visual accuracy for relative depth is maximal, suggesting that motor coordinates are constitutive for the calculation of object relations. A visual discrimination task was presented before and after pointing adaptation to estimate the putative effect of motor signals on visual space. Measuring purely visual adaptation aftereffects with a psychophysical task requires presenting a probe in an adapted region and a comparison stimulus in a non-adapted region of the visual field. We used a dual adaptation method where pointing to the right was followed by wrong visual feedback about the movement and pointing to the left by correct feedback. It has been shown that humans are able to adapt identical movements to two (or more) perturbations simultaneously as long as a contextual cue differentiates between the two (Osu et al., 2004; Nozaki et al., 2006; Howard et al., 2008; Shelhamer et al., 2005; Imamizu et al., 2007; Choi et al., 2008; Gandolfo et al. 1996; Hirashima and Nozaki 2012; Howard et al. 2012; Sarwary et al. 2013). Ghahramani et al. (1996) demonstrated that it is possible to produce a dual adaption field where rightward movements are adapted in one direction and leftward movements in the opposite direction.

## Methods

### Experiment 1

#### Participants

Forty-five subjects (28 females, ages 18-47, average 25, 4 left-handed), including the second author, participated in the experiments. In separate sessions, we tested two different adaptation directions. Participants were randomly assigned to one of these two session types. In shortened reach adaptation, 23 subjects participated and in extended reach 22. All participants had normal or corrected-to normal vision. All participants gave their written informed consent prior to participation and subsequently received either monetary compensation or course credits. All experiments were approved by the ethics committee of the Faculty of Mathematics and Natural Sciences of the Heinrich-Heine-University Düsseldorf, Germany and the study procedures were in line with the declaration of Helsinki.

#### Apparatus and stimuli

Participants were sitting in a chair wearing a head-mounted display (HMD) and holding a VR controller in their right hand. Stimuli were delivered by an Intel i7-based PC (Intel, Santa Clara, U.S.) with an NVIDIA GTX 1080 connected to an HTC Vive HMD (HTC Corporation, Taoyuan, Taiwan). The HMD presents stimuli on two low-persistence organic light-emitting diode (OLED) displays with a resolution of 1,080 × 1,200 pixels per eye and a refresh rate of 90Hz. Additionally, participants received a Vive motion-controller for their right hand. The virtual environment was rendered using a custom-made program created in the Unity game engine, version 2019.1.8f1 (Unity Technologies, San Francisco, U.S.). Head and hand movements were tracked via the HMD and controller using the standard SteamVR tracking system. According to previous research (Niehorster et al., 2017, Verdelet et al. 2019), this tracking system provides a robust tracking of head and hand positions with a 360° coverage provided tracking loss is prevented. In the present study, participants always responded to stimuli in front of the and we did not need the full coverage around the participant. Hence, in order to minimize the change of occlusions of the HMD or controller and thereby avoiding tracking loss, our setup had both base stations facing the participant. Throughout the experiment, participants held the controller with an outstretched index finger placed on top of the controller with the fingertip matching the tracking origin of the controller as close as possible.

#### Target positions

Stimuli were presented in an empty mid-gray space maximally reducing monocular cues. In order to ensure that the locations of the stimuli relative to the observer were the same across participants, all stimuli were positioned relative to the location of the HMD at the start of the experiment. Participants were instructed to the stay in the same position throughout the experiment (see Figure 1A). A visual and acoustic error signal appeared when the participants left their predetermined head position.

**Figure 1.**
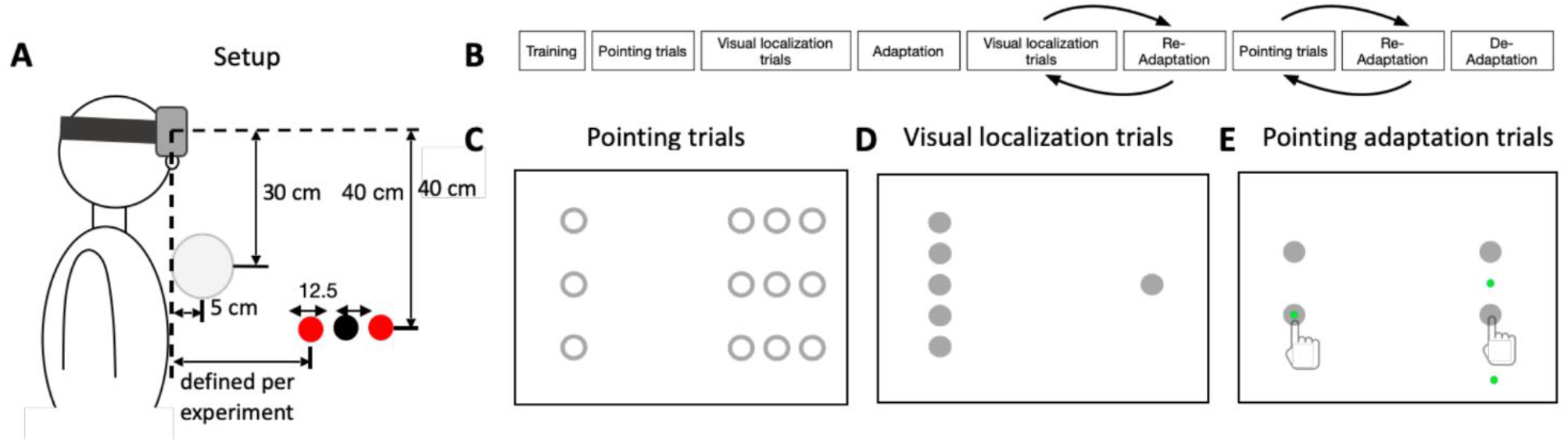
**A** Side view of the possible stimulus positions relative to the observer along the z-axis in pointing trials of Experiment 1 and 2. Only one target was shown per trial. **B** Trial structure for Experiments 1 and 2. **C** Locations of all pointing targets in Experiment 1. In Experiment 2 only 2 of these targets were presented. **D** Locations of all visual localization targets in Experiment 1. In Experiment 2 a different procedure was used (see methods section for more information). **E** Locations of all pointing adaptation targets in Experiment 1 and 2. On the left side the feedback was accurate, on the right it was distorted such that to hit the target observer should either point closer than the target (extended reach), or further than the target (shortened reach). Green dot shows the pointing feedback resulting from respective hand positions.

A white sphere (diameter: 10 cm) was placed centrally 5 cm in front of their chest. This sphere served as a starting position where participants had to place their right hand between trials. Pointing targets were colored red and visual targets black (diameter: 3 cm). The distance between the participant and the pointing targets was adjusted according to the arm length of the participants. This was done in order to match targets around the natural grasping distance. To this end, participants extended their hand forward as far as they could. In the experiments, targets were presented such that the farthest targets in z-direction were 15 cm closer to the participant than the final position of the hand when the arm was fully extended. In pre-and post-adaptation pointing trials,12 targets (nine on the right side, three on the left side) were presented in total, one per trial. On both sides, the distance between pointing targets in z-direction was set to 12.5 cm. On the right side, the distance between pointing targets in x-direction was set to 7 cm.

In adaptation trials, a target appeared in one of four possible target locations (with two on the left and two on the right side). The four target positions were spaced 15 cm in z-direction and 30 cm in x-direction. In adaptation, observers made 40 repetitions per target location. The feedback on the left side was always veridical, and on the right side distorted to shorten or extend participant’s reach, depending on the session’s condition (shortening or extending the reach). The 10 cm distortion was introduced gradually over three quarters of the whole amount of trials in order to minimize conscious adjustments of pointing (recalibration).

In visual localization trials, observers saw two targets in front of them, a static reference and a probe whose position in z-direction could be controlled with their thumb using the controller’s trackpad. In half of the trials, the reference sphere was presented on the left side and the probe on the right and vice versa. In total 10 probe positions were tested (five on each side), one per trial. The distance between the probe positions in z-direction was set to 6.25 cm. Participants were instructed to adjust the position of probe along the z-axis until it matches depth of the reference stimulus. Each position of the reference was repeated 5 times.

Targets in de-adaptation trials were the same as in adaptation trials. However, in de-adaptation trials, participants always received veridical feedback.

#### Trial structure Experiment 1

Before the experiments started, the participants were trained for 40 trials to execute a pointing movement to a peripherally flashed target with their right index-finger. In each trial, a single red sphere appeared for 500 ms in front of the participant. Participants were instructed to point to the sphere reaching it with their index finger. The pointing was registered when the tracked finger reached the position of the target which was signaled to the participant via a short vibration of the controller. During the training, the participants received an accurate visual feedback of their landing position in form of a green sphere (1 cm in diameter).

The experiments consisted of five phases (see Figure 1B). In 120 pointing trials, participants had to point to 1 of 12 possible target positions, nine on the right and three on the left (see Figure 1C). Each position was tested 10 times. In all trials except pre- and post-adaptation pointing trials, participants also received a visual feedback of their landing position, accurate or distorted depending on trial type and pointing side. When participants returned their hand to the starting position, another short vibration was emitted, and the next trial started after a fixed inter-trial interval of 1000 ms.

Then, a block of 50 visual localization trials followed. In a visual localization trial, observers saw two black spheres. One of them served as reference and the other as probe whose position along the z-axis could be controlled via the trackpad of the controller. The observers were asked to adjust the position of the probe sphere until it matched the location of the reference sphere in z-direction.

In 160 adaptation trials (see Figure 1E), participants received wrong feedback about their terminal pointing location. In shortened reach sessions, the feedback appeared closer in depth relative to the actual pointing position. In extended reach sessions the feedback was shown farther in depth. The displacement of the feedback shown on the display was increased gradually across trials until it reached the maximum value of 10 cm. In an adaptation trial, a target appeared in one of four possible positions, two targets positions were in the left half of the display and two on the right. After pointing to the targets on the left side, participants always received veridical visual feedback about their terminal pointing location, whereas in adaptation trials, pointing to targets on the right side was followed by systematically shifted feedback.

After the adaptation block, participants performed four blocks of the visual localization again, each block consisting of 16 trials, except the last block, which contained only two trials, resulting in 50 visual localization trials in total. Between each block, participants had to perform a block of eight re-adaptation trials. Then seven blocks of 16 pointing trials were conducted, again was each block separated by a block of eight re-adaptation trials. Following the last re-adaptation block, a last pointing block of eight trials was performed, resulting in 120 pointing trials in total.

Finally, 20 de-adaptation trials were conducted, without any distorted feedback.

Experiment 1 contained 640 trials and took on average around 37 minutes.

### Experiment 2

#### Participants

Fifty-one participants (31 females, ages 18-47, average 25, 2 left-handed), including two of the authors, participated in Experiment 2. All participants had normal or corrected-to normal vision. Each subject participated in two adaptation session (shortened reach and extended reach adaptation). All participants gave their written informed consent prior to participation and subsequently received either monetary compensation or course credits.

#### Target positions

Target positions in Experiment 2 were the same during adaptation as in Experiment 1 and differed only in pre-and post-adaptation pointing and visual localization trials.

In pointing trials, the number of possible target positions was reduced to two (one on the left and one on the right). The positions corresponded to the central target position in the left group of targets shown in Figure 1C and the central target position of the group of targets shown on the right. The closer pointing target positions were positioned 30 cm from the headset for shortened reach adaptation sessions and 40 cm for extended reach adaptation sessions in order to attend for the distances that observers needed to reach. With this change we aimed to have the final adapted movements comparable in shortened reach and extended reach adaptation.

In the visual localization trials, two stimuli were presented (one on the left side and one on the right) with an inter-stimulus interval of 250 ms. The stimulus on the right side was the probe stimulus that way always presented in the same location which corresponded to the central location in the right side (see Figure 1C). On the left side, the reference stimulus was presented in one of seven possible, equiprobable and equidistant (2 cm) locations.

#### Trial structure experiment 2

The procedure of Experiment 2 followed the trial structure of Experiment 1 (Figure 1B). After the training block, participants performed 20 pointing trials without visual feedback towards two target locations with 10 repetitions per location.

Then, a block of 98 visual localization trials was presented. A trial started with the presentation of a fixation point for 1000 ms, to which participants had to direct their gaze. Then, the left black sphere was flashed at one of the seven positions on the left for 10 ms, and after 200 ms delay, the right sphere was flashed also for 10 ms. Participants were asked to indicate which sphere was closer to them (their body) in depth by moving controller to the left or right side of the starting position. Recording of their feedback was indicated by a vibration, as well as the following return to the starting position. A psychometric function was measured for two probe locations. For each psychometric function, seven different comparison stimuli were presented, which each were tested seven times.

Following adaptation, participants performed visual and blind testing blocks again, mixed with re-adaptation blocks in order to maintain the adaptation effect. For each 10 testing trials, participants had eight re-adaptation trials (two repetitions per adaptation target position). Finally, participants made five de-adaptation trials per target in which they received an accurate pointing feedback on both sides. Target presentations in the pointing and the visual trials were randomized. Altogether, the experiment contained 544 trials and took around half an hour to complete.

Each participant performed two sessions (shortened reach and extended reach adaptation), separated by at least a few hours, and for most participants (37 of 41) were separated by a day and more.

#### Data analysis

The free statistical software R (R Foundation for Statistical Computing, Vienna, Austria; www.r-project.org) and RStudio (RStudio Team, 2020) were used to analyze the behavioral data. For statistical analysis, a non-parametric repeated measures ANOVA was calculated, using the Aligned Rank Transform (Wobbrock et al., 2011). Significance was determined by applying the Kenward-Roger approximation to estimate p-values, a procedure that has been shown to produce acceptable Type 1 error rates (Luke, 2017).

Participants were instructed to point to the targets from below. We determined the terminal pointing coordinates in the x- and the z-direction for a given target position once they crossed the table level in the y-direction. We included all trials in the analysis, where the pointing movement did not start before or too late after target onset. To this end, we defined pointing trajectories as valid if they lasted longer as 100 ms and were shorter than the mean plus two standard deviations of the path length for the respective observer. In Experiment 1, we included 95,4% in the shortened reach adaptation sessions and 94,6% in the extended reach adaptation sessions.

For the analysis of Experiment 2, we excluded again trials from pre- and post-adaptation pointing trials, using the same criteria as in experiment 1. We included 94.5% of the data in the shortened reach adaptation sessions and 96,4% in the extended reach adaptation sessions.

## Results Experiment 1

We first checked the pointing performance of the 120 pre-adaptation pointing trials in which participants received no visual feedback about their terminal pointing position. In each trial, participants pointed to a target in one of 12 possible positions. Three targets positions where on the left side and nine on the right. Targets presented on the right side served as probe targets to investigate generalization of pointing adaptation. For the probe targets, we analyzed pointing errors in the z-direction by averaging for each participant across all three targets in the x-direction. Figure 2A shows average pre-adaptation pointing errors from sessions with shortened reach adaptation. Positive errors for the close targets indicate that participants overshot the physical target location with their pointing. The opposite holds true for the far targets. These were undershot by participants. A non-parametric repeated measures ANOVA revealed a significant effect of target depth on pointing error size (F(2, 43) = 20.43, p = 5.79 × 10^−7^). Average pre-adaptation pointing errors from sessions with extended reach adaptation are shown in Figure 2B. Similarly, subjects overshoot the close and undershot the far targets. Pointing errors size differed significantly between targets in depth (F(2, 41) = 8.75, p = 0.0007).

**Figure 2.**
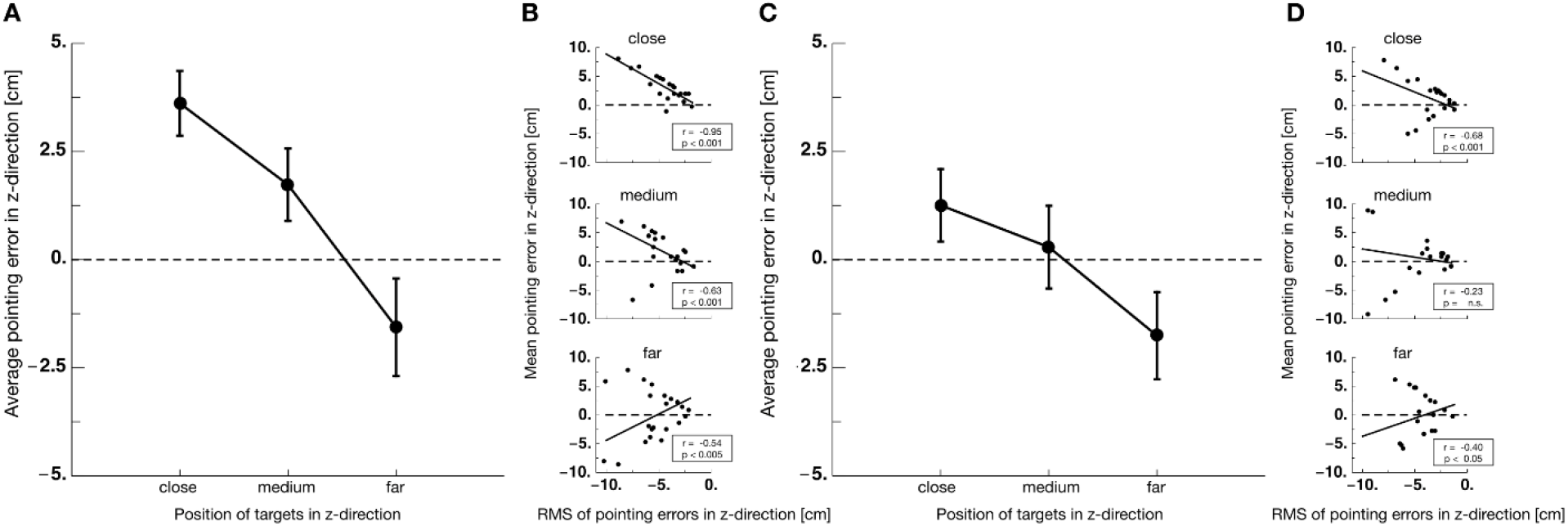
**A** Pointing errors for all three target distances in z-direction averaged within participants across all targets in x-direction and averaged across observers. Data derive from shortened reach adaptation sessions. Error bars represents S.E.M. across observers. **B** Mean pointing errors in z-direction against root mean square error of pointing for all observers in shortened reach adaptation sessions. The black line indicates the linear regression. **C** Pointing errors for all three target distances in z-direction averaged within participants across all targets in x-direction and averaged across observers. Data derive from extended reach adaptation sessions. Error bars represents S.E.M. across observers. **D** Mean pointing errors in z-direction against root mean square error of pointing for all observers in extended reach adaptation sessions. The black line indicates the linear regression.

After the pre-adaptation trials, 50 pre-adaptation visual localization trials were presented, which will be analyzed below, together with the post-adaptation visual localization trials. Then, in 160 adaptation trials (40 trials of each of the two targets on the left and on the right side), participants received visual feedback about their terminal pointing position once their pointing movement was finished. The location of the feedback was gradually distorted over trials, i.e. it was displayed shifted from the physical terminal position. The distortion reached a maximum of 10 cm. Feedback was only distorted for the targets on the right but not for those on the left side. Figure 3A shows average pointing performance in the shortened reach adaptation trials for each of the two targets on the left side. Although no distorted feedback was provided for pointing to targets on the left side, one can see that across trials, participants pointed closer to their bodies, thus undershooting the physical target location. In shortened reach adaptation sessions, feedback was presented closer to the body of the observers. In attempting to reduce the error between their desired pointing location and the visual feedback, participants point closer to their bodies. As feedback was distorted only on the right side, adaptation likely overlapped to the left side, generating this small trend of adaptation seen in Figure 3A. Terminal positions for pointing to targets on the right side is shown in Figure 3B. For both targets a strong adaptive shift in pointing terminal positions was generated by the distorted feedback. In sessions with feedback that suggested an extended reach, participants should point farther in depth, trying to reduce the error between their terminal position and the feedback location. Similar as in shortened reach sessions, adaptation was very weak on the left side (see Figure 3C). Only a small amount of adaptation over the course of trials is visible. However, on the right side a strong adaptation magnitude can be seen (see Figure 3D).

**Figure 3.**
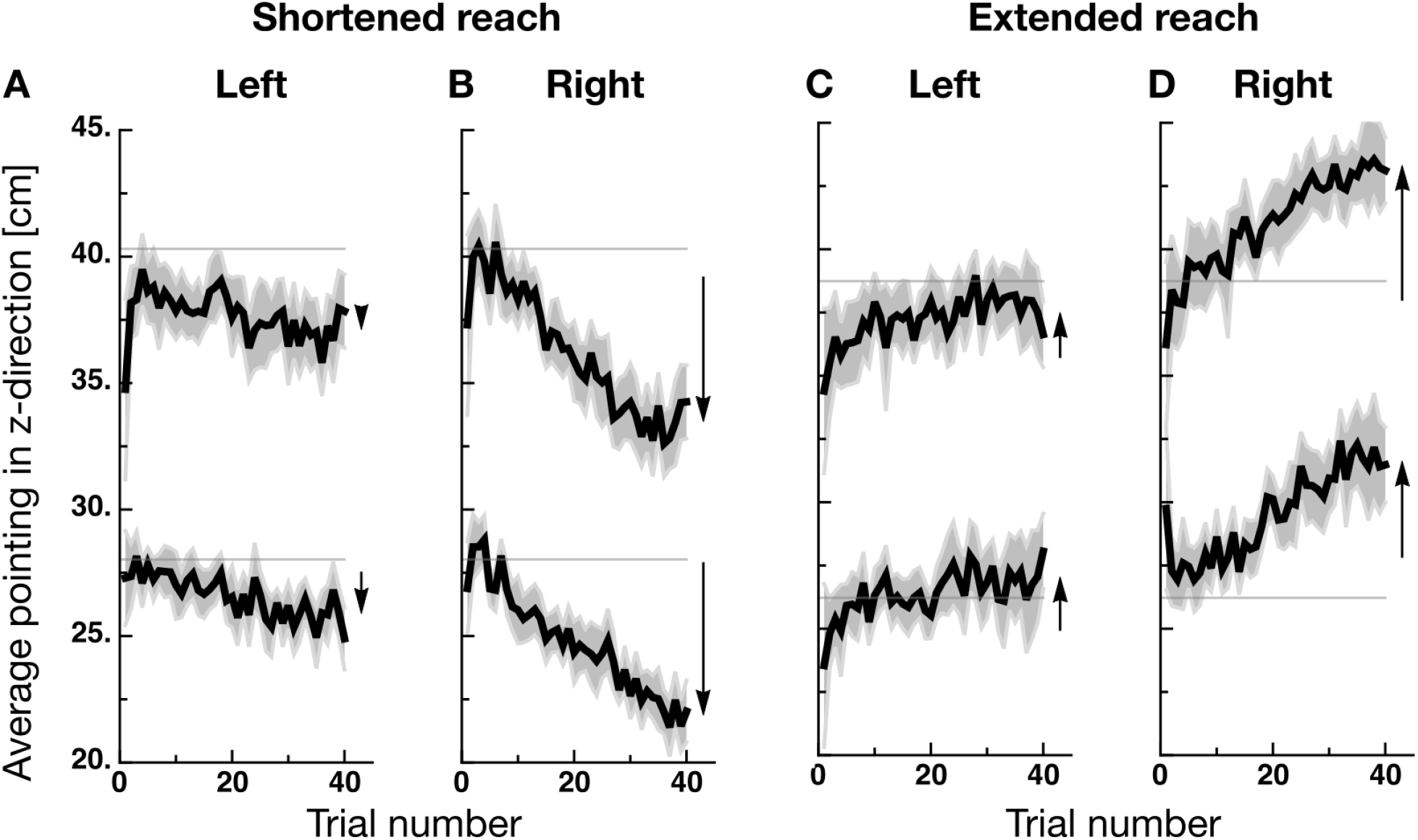
**A-D** Adaptation curves averaged over participants, one curve for each target position. Horizontal grey lines mark the target position. Black arrows on the right of the curves connect the average values of first and last 5 trials of respective curve. Vertical dashed line represents the end of the distortion adjustment. Error bands represent 95% confidence intervals.

In order to estimate whether adaptation took place at the movement planning stage, we analyzed the peak velocities of the movement trajectories. Figure 4A shows average peak velocities from shortened reach adaptation sessions for the left and the right side from pre-adaptation pointing trials (shown in white) and post-adaptation pointing trials (shown in black). Peak velocities of pointing movements to targets on the left were virtually identical. However, peak velocities of pointing movements to targets on the right side were higher before than after adaptation. This difference is consistent with the smaller movements, i.e. pointing closer to the body, that was performed after shortened reach adaptation. A non-parametric repeated measures ANOVA revealed a significant main effect for pointing direction (left / right: F(1, 22) = 33.23, p = 8.45 × 10^−8^) and for adaptation phase (pre/ post: F(1, 22) = 11.73, p = 0.002). In extended reach adaptation session, peak velocities did not differ much on the left side. On the right side, peak velocities were higher after adaptation than before adaptation, consistent with movements that were farther in the z-direction. A non-parametric repeated measures ANOVA revealed a significant main effect for pointing direction (left / right: F(1, 21) = 7.03, p = 0.015) and for adaptation phase (pre/ post: F(1, 21) = 4.99, p = 0.037) and a significant interaction effect (F(1, 21) = 29.13, p = 2.35 × 10^−5^).

**Figure 4.**
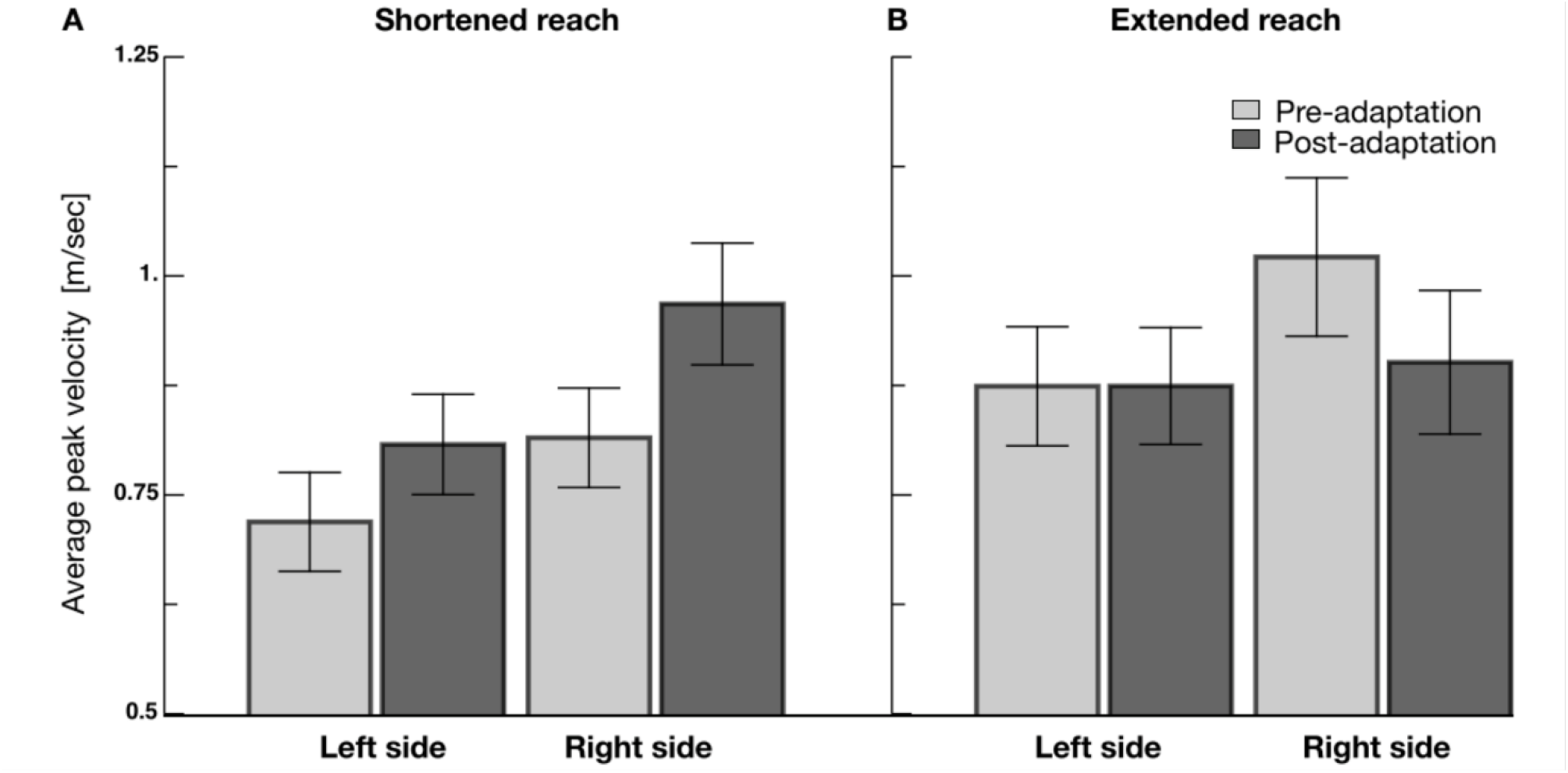
**A** Average peak velocity of pointing movements from pre-adaptation (bright gray) and post-adaptation trials (dark gray) in the shortened reach adaptation sessions. Error bars represent S.E.M.**B** Average peak velocity of pointing movements from pre-adaptation (white bars) and post-adaptation trials (black bars) in the extended reach adaptation sessions. Error bars represent S.E.M.

After adaptation trials were finished, post-adaptation pointing trials alternated with re-adaptation trials. We defined pointing aftereffects as the difference between pointing before and after adaptation. Figure 5A shows aftereffect magnitudes for each of the 12 probe targets as arrows. The starting point of each arrow is given by the average pre-adaptation pointing location and the tip of the arrow represents the average post-adaptation pointing location. Circles show the physical target positions. For all arrows on the right side it can be seen that participants pointed farther in z-direction, indicating that shortened reach adaptation successfully increased pointing movements in depth. One can also see that before and after adaptation, pointing to targets close to the body overshot the physical target position while pointing for the targets farthest in z-direction were closer to the physical target locations. The tendency to overshoot the close targets was already analyzed for pre-adaptation trials (see Figure 2). Due to the adaptation direction in shortened reach sessions, this tendency increased after adaptation. No systematic pointing aftereffects were found for pointing to targets on the left side. Average results from extended reach sessions are shown in Figure 5B. As for shortened reach adaptation, aftereffects were consistent for all targets on the right side. For all nine targets pointing movements were decreased in z-direction. Again, no systematic aftereffects were found for pointing to targets on the left side. When comparing aftereffect magnitudes on the right side for shortened and extended reach adaptation, one can see that for shortened reach adaptation aftereffects are stronger for the closer targets than for the farthest targets in z-direction. In extended reach adaptation, the opposite holds true. Pointing to the farthest targets shows the strongest adaptation aftereffects. For statistical analysis we used pointing errors, defined as the difference between each pointing terminal position and the physical target location. For shortened reach adaptation, a non-parametric repeated measures ANOVA with the factors target position in x-direction (three steps), target position in z-direction (three steps) and adaptation phase (pre / post) revealed a significant main effect target position in z-direction (F(2,44) = 31.63, p = 3.06 × 10^−9^), a significant main effect adaptation phase (F(1,22) = 23.86, p = 6.97 × 10^−5^). The main effect for target position in x-direction was not significant (F(2,44) = 0.08, p = 0.919). The interaction between adaptation phase and target position in x-direction was significant (F(2,44) = 3.57, p = 0.037). These results confirm that adaptation changed pointing terminal positions and that adaptation magnitude was modulated by the target position in z-direction. Furthermore, the interaction also indicated a modulation of the adaptation magnitude by the target location in x-direction. Neither the interaction between target position in x-direction and target position in z-direction (F(4.88) = 2.28, p = 0.067) nor the interaction between target position in z-direction and adaptation phase (F(2,44) = 2.55, p = 0.089) were significant.

**Figure 5.**
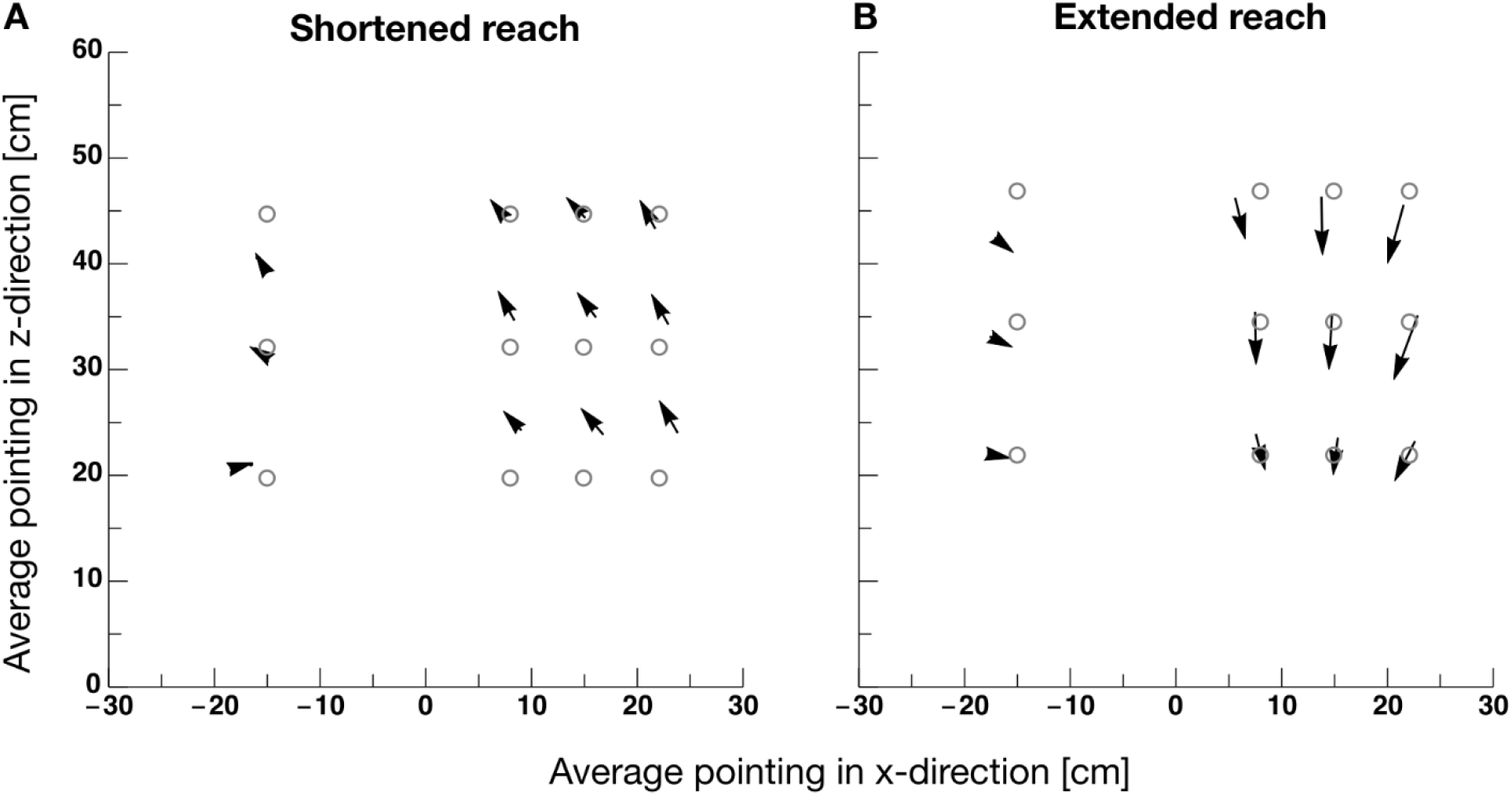
**A** Average pointing aftereffects in shortened reach sessions shown as arrows. The x-direction of the pointing trajectory is shown on the abscissa and the z-direction on the ordinate. Gray circles mark the positions of targets. The starting point of the arrows indicate the average pre-adaptation pointing location and the tips the average post-adaptation pointing location. **B** Average pointing aftereffects in extended reach sessions. Same conventions as in **A**.

For extended reach adaptation, a non-parametric repeated measures ANOVA with the factors target position in x-direction (three steps), target position in z-direction (three steps) and adaptation phase (pre / post) revealed a significant main effect target position in x-direction (F(2,42) = 17.73, p = 2.62 × 10^−6^), a significant main effect target position in z-direction (F(2,42) = 13.47, p = 3.02 × 10^−5^), a significant main effect adaptation phase (F(1,21) = 43.07, p = 1.68 × 10^−6^) and a significant interaction effect adaptation phase and target position in x-direction (F(2,42) = 4.65, p = 0.014) a significant interaction effect adaptation phase and target position in z-direction (F(2,42) = 9.53, p p = 3.86 × 10^− 4^). The interaction between target position in x-direction and target position in z-direction was not significant (F(4,84) = 1.34, p = 0.261). Similarly, the interaction between target position in x-direction, target position in z-direction and adaptation phase was not significant (F(4,84) = 2.44, p = 0.053). As for shortened reach adaptation, these results confirm that adaptation successfully change pointing behavior and that adaptation magnitude depended on x-direction and z-direction of then targets.

Our analysis of pointing errors in pre-adaptation pointing trials revealed a dependence of errors on target z-direction (see Figure 2).

We also wondered whether the pre-adaptation bias determined adaptation magnitude. In shortened reach adaptation for instance, one could expect that participants who undershoot the targets would adapt stronger as the pre-adaptation bias and the undershoot would add up. Similarly, for extended reach adaptation, overshooting physical target positions and visual feedback should add up and lead to higher adaptation magnitudes. To investigate this putative dependency, we determined average pointing errors in z-direction for each participant and correlated these with pointing aftereffects across all subjects. Figure 6A shows a significant negative correlation between pre-adaptation errors and pointing aftereffects derived from shorted reach adaptation sessions. This correlation indicates that the more subjects undershoot before adaptation, the higher is adaptation magnitude. Figure 6B shows a significant positive correlation for extended reach adaptation sessions. The direction of these correlations is consistent with the idea that the natural pointing error - measured before adaptation - increased the visual feedback distortion displayed after pointing and thereby enlarged adaptation magnitude.

**Figure 6.**
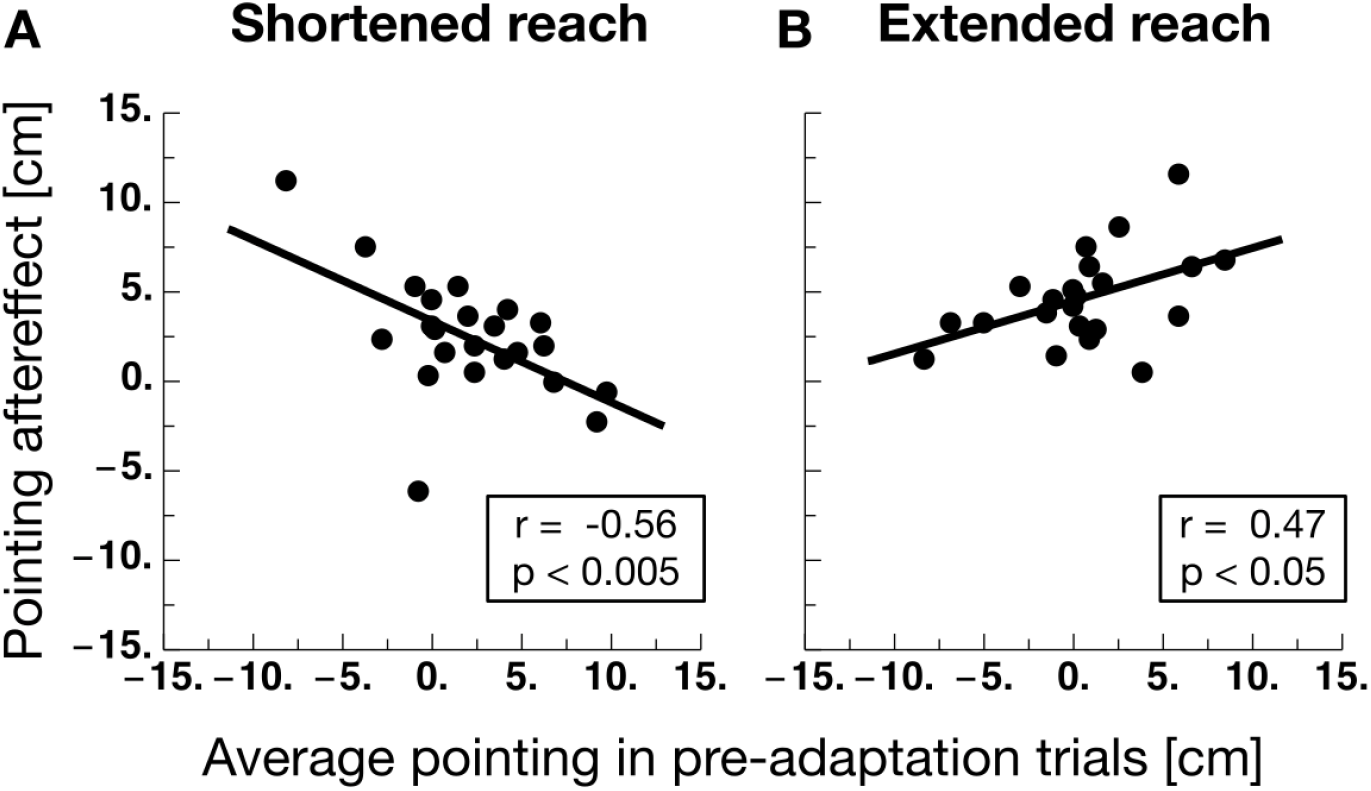
**A** Pointing aftereffects magnitude against average pointing errors from pre-adaptation trials for each participant from shortened reach adaptation sessions. The black line represents the linear regression. **B** Pointing aftereffects magnitude against average pointing errors from pre-adaptation trials for each participant from extended reach adaptation sessions. Same conventions as in **A**.

We measured visual localization by asking participants to match the position of a visual reference by adjusting the position of a visual stimulus presented on the opposite side of the visual field. Visual localization was measured before and after adaptation. Figure 7A shows visual localization aftereffects for all 10 target locations from shorted reach adaptation sessions. As for pointing, visual aftereffects were defined as the difference between pre- and post-adaptation and are represented by the lengths of the arrows. For all positions except one, after adaptation targets were localized to be farther in z-direction. A non-parametric repeated measures ANOVA with the factors positions (five), adaptation phase (pre / post) and adaptation region (left / right) revealed a significant main effect for the factor positions (F(4, 88) = 14435, p = 2.22 × 10^−16^) and a significant main effect for the factor adaptation phase (F(1, 22) = 4.55, p = 0.044). The factor adaptation region did not result in a significant effect (F(1,22) = 0.63, p = 0.437). The interaction between positions and adaptation phase was significant (F(4, 88) = 3.55, p = 0.009). Neither the interaction between positions and adaptation region (F(4,88) = 1.92, p = 0.114) nor the interaction between adaptation phase and adaptation region (F(1,22) = 1.33, p = 0.262) were significant. Similarly, the interaction between positions, adaptation phase and adaptation region was not significant (F(4,88) = 0.48, p = 0.745).

**Figure 7.**
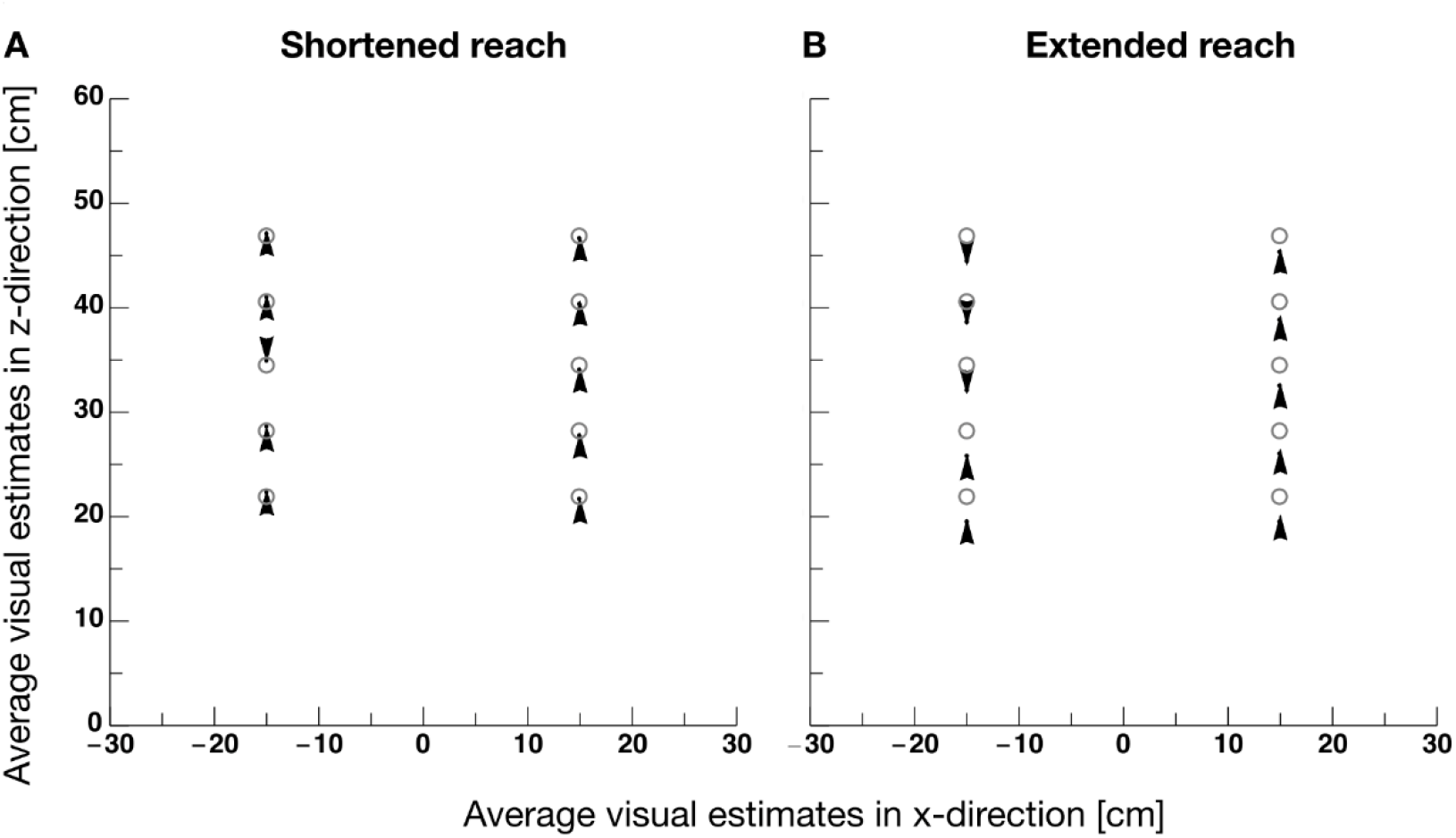
**A** Average localization errors from localization trials in shortened reach adaptation sessions shown as arrows. The starting point of the arrows indicate the average pre-adaptation localization location and the tips the average post-adaptation localization location. Error bars represent S.E.M. **B** Average localization errors from localization trials in shortened reach adaptation sessions shown as arrows. Same conventions as in **A**.

Aftereffects measured in extended reach adaptation sessions are shown in Figure 7B. Visual localization of targets on the right side is generally shifted into the z-direction after adaptation visual localization of targets on the left side is not systematically shifted for all positions. A non-parametric repeated measures ANOVA revealed a significant main effect for the factor positions (F(4, 84) = 1.30, p = 2.0 × 10^−16^), a significant main effect for the factor adaptation phase (F(1, 21) = 6.06, p = 0.02). The main effect for the factor adaptation side was not significant (F(1,21) = 0.16, p = 0.696). The analysis revealed no significant interactions between positions and adaptation phase (F(4,84) = 0.32, p = 0.864), positions and adaptation side (F(4.84) = 0.37, p = 0.831) or adaptation side and adaptation phase (F(1,21) = 0.46, P = 0.504). The interaction between positions, adaptation phase and adaptation side was not significant (F(4,84) = 0.49, p = 0.75).

The absence of a significant interaction effect for both adaptation directions leaves it open whether the pointing shift in z-direction was triggered by adaptation. Distorted visual feedback was only provided on the right side. If adaptation were responsible for the pointing shift in z-direction, one would have expected a stronger shift on the right than on the left side. In order to investigate that hypothesis with a more sensitive visual localization task, we conducted a second experiment.

## Results Experiment 2

Experiment 2 aimed at testing visual localization after pointing adaptation with a more sensitive method. We tested pointing adaptation with only two target positions. In trials of the visual task, we flashed a target on the unadapted left side and one on the adapted right side in order to implement a psychometric spatial discrimination task. The trial structure was identical to Experiment 1 (see Figure 1B), only the number of trials was changed for certain blocks.

Since we measured only two target positions before and after adaptation, 20 trials were tested in each the pre- and the post-adaptation pointing blocks. The adaptation block contained 160 trials as in Experiment 1 with pointing to 4 different target locations. Pointing aftereffects are shown in Figure 8. In shortened reach adaptation sessions, pointing to the target on the left side, where visual feedback was veridical during adaptation, remained unchanged after adaptation. However, pointing to the target on the right side was shifted in z-direction after adaptation. A non-parametric repeated measures ANOVA with the factors positions (left / right) and adaptation phase (before / after) revealed a significant main effect for the factor positions (F(1, 50) = 245.78, p = 2.22 × 10^−16^), a significant main effect for the factor adaptation phase (F(1, 50) = 100.64, p = 1.44 × 10^−13^) and a significant interaction between positions and adaptation phase (F(1, 50) = 141.57, p = 3.38 × 10^−16^). These results confirm that distorted visual feedback successfully adapted pointing behavior. The same hold true for extended reach sessions, except that then direction of the aftereffect for pointing on the right side reversed its direction. A non-parametric repeated measures ANOVA revealed a significant main effect for the factor positions (F(1, 50) = 84.17, p = 2.68 × 10^−12^), a significant main effect for the factor adaptation phase (F(1, 50) = 8.66, p = 0.005) and a significant interaction between positions and adaptation phase (F(1, 50) = 81.23, p = 4.71 × 10^−12^). The size and direction of the aftereffects for both, the shortened reach and the extended reach adaptation, replicate the findings from Experiment 1.

**Figure 8.**
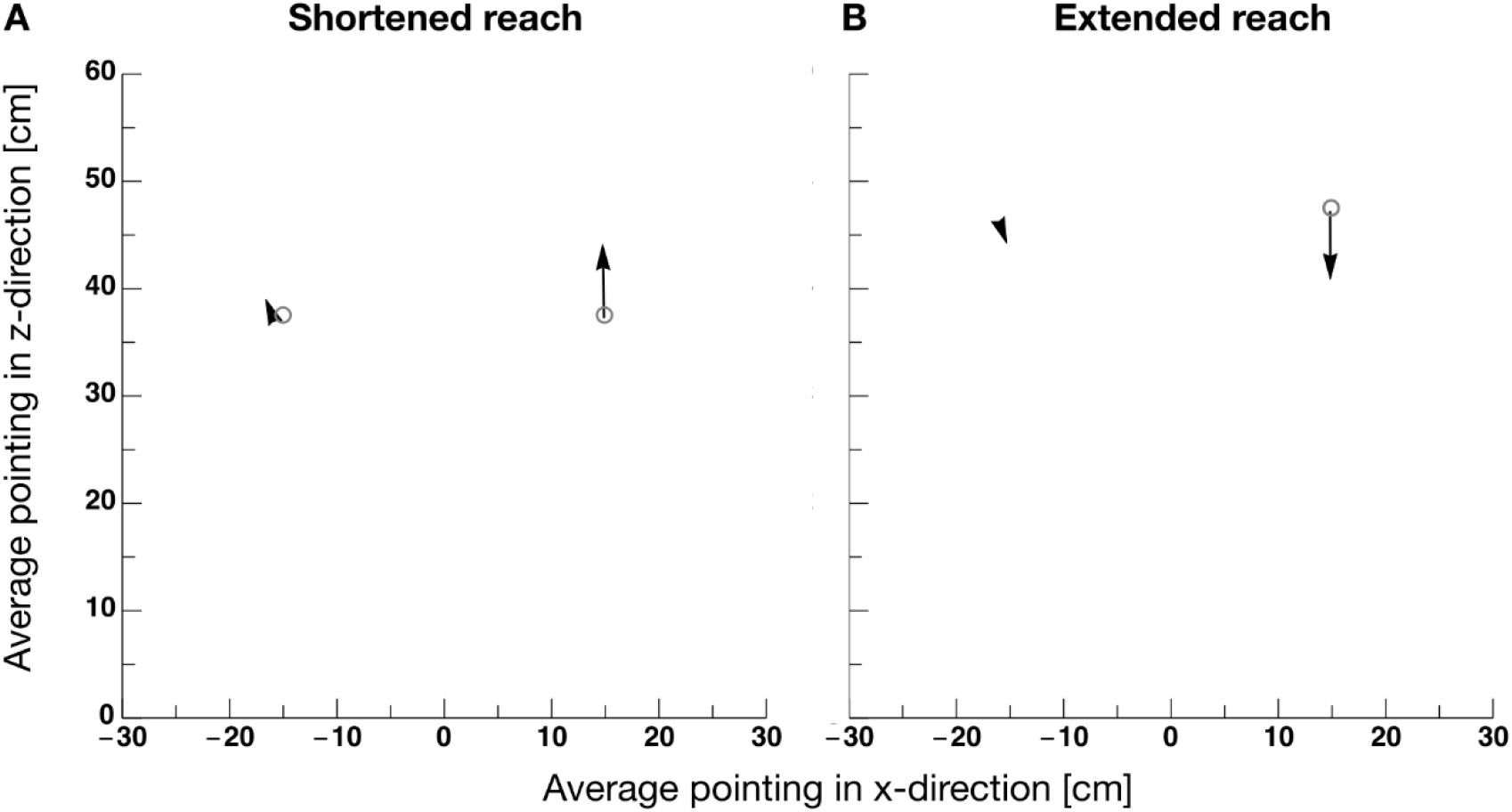
**A** Average pointing aftereffects in shortened reach sessions shown as arrows. The x-direction of the pointing trajectory is shown on the abscissa and the z-direction on the ordinate. Gray circles mark the positions of targets. The starting point of the arrows indicate the average pre-adaptation pointing location and the tips the average post-adaptation pointing location. **B** Average pointing aftereffects in extended reach sessions. Same conventions as in **A**.

In 98 visual discrimination trials, tested before and after adaptation, participants estimated which target, the one on the left or the one on the right, was located farther in z-direction. Figure 9A shows example psychometric functions from two observers from shortened reach adaptation sessions. Psychometric functions measured before adaptation are shown in gray and those measured after adaptation are shown in black. The slope of the curves reveals that participants were well able to perform the discrimination task. However, there is no difference in localization before and after adaptation for these two participants. On average, localization was almost veridical with no difference before and after adaptation (see Figure 9A, paired t-test, t(50) = −1.29, p = 0.19). Figure 9C shows example psychometric functions from two observers from extended reach adaptation sessions. The results were almost identical to those of shortened reach adaptation. Localization was nearly veridical and did not differ between adaptation states. The same holds true for the average performance, as can be seen in Figure 9D (paired t-test, t(50) = −0.67, p = 0.50).

**Figure 9.**
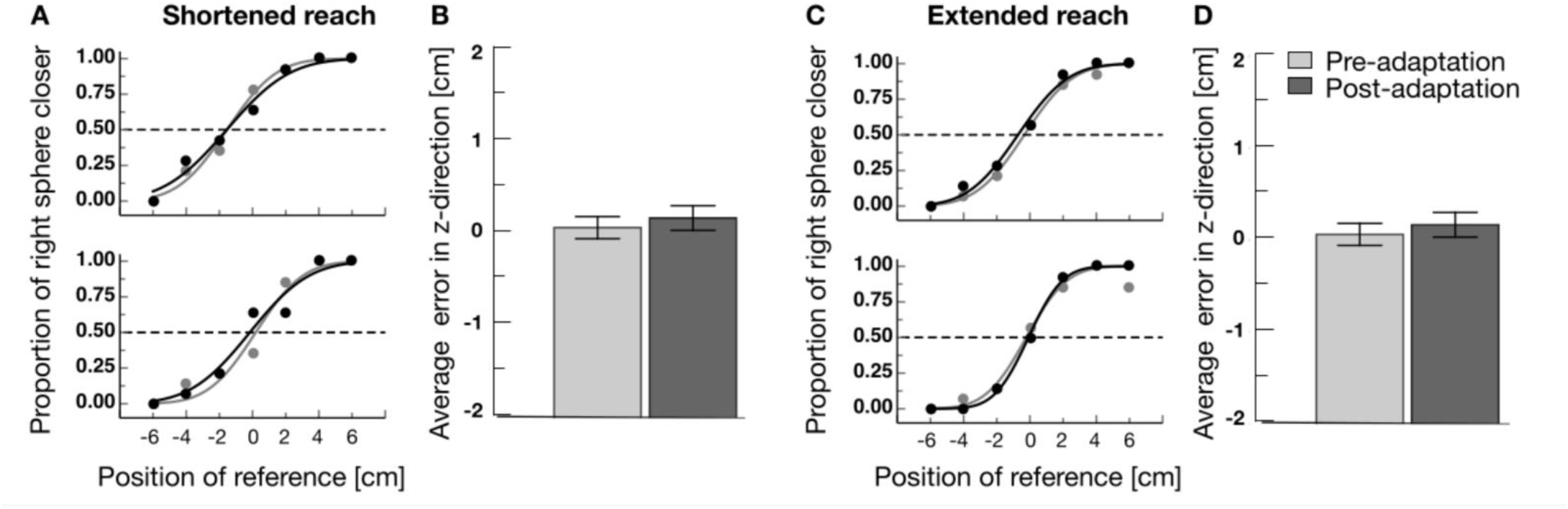
**A** Example psychometric functions from shortened reach adaptation sessions from before (gray curves) and after adaptation (black curves). **B** Average localization errors for pre-adaptation (bright gray bar) and post-adaptation trials from shortened reach adaptation sessions. **C** Example psychometric functions from extended reach adaptation sessions from before (gray curves) and after adaptation (black curves). **D** Average localization errors for pre-adaptation (dark gray bar) and post-adaptation trials from extended reach adaptation sessions.

## Discussion

In this study we asked whether the distortions in visual 3D space, that are commonly observed in human perception, are compensated for pointing by involving prior knowledge about the natural grasping distance. To this end, we investigated the spatial generalization of pointing adaptation. We first looked at pointing errors before adaptation. We found that like the visual distortions, participants estimated close targets to be further in depth and far targets to be nearer than they really are. This bias might have two origins: On the one hand, participants might produce these motor errors because they are guided by distorted space perception, on the other hand, movement planning might be biased towards the “natural grasp distance”. In the pre-adaptation data, we already found correlative evidence for the latter hypothesis. Those subjects with the highest variable errors showed the strongest constant errors in their pointing. We used pointing adaptation to find causal evidence that might distinguish between both hypotheses. First, we applied visual pointing feedback that suggested a shortened reach. This artificial error led participants to extend their movements further into depth. We found a significant difference in adaptation strength in depth. Adaptation was stronger for targets close to the body and weakest for targets farthest in depth. This result is in principle consistent with adaptation following the visual distortion. Since vision is overestimated for targets being closer to the body, the distorted feedback should be overestimated for close targets and therefore induce stronger adaptation. Under this hypothesis the same modulation of adaptation magnitude as a function of distance in depth would be expected for extended reach adaptation. By contrast, the involvement of a “natural grasping distance prior" implies the opposite hypothesis. In that perspective, adaptation should be strongest when leading towards the “natural grasping distance” and weakest when leading away from it. The results from extended reach adaptation are clearly in favor of the latter view. Adaptation was weakest for close targets and strongest for targets farthest in depth. In summary, adaptation generalization in depth for both adaptation directions are biased by an urge towards the middle position that can also be described as the “natural grasping distance”.

Importantly, we also found significant negative correlations between pre-adaptation motor error and pointing adaptation magnitude. This result provides a hint about the multi-sensory discrepancy that led to adaptation. Our distorted visual pointing feedback induces a discrepancy between the felt and the seen terminal pointing location. Adaptation induced by this discrepancy is consistent with the negative correlation that we found. Those participants who overshoot strongest because they see targets furthest in depth will also see the terminal visual pointing feedback further in depth. The terminal visual pointing feedback position is displaced by a constant amount relative to the terminal pointing location. Thus, participants who see stimuli further in depth, receive an error signal with a smaller size than other participants. A smaller error signal will lead to a reduced adaptation magnitude.

In two experiments, we tested visual adaptation in a two alternative forced choice task and in an adjustment task before and after adaptation. Both tasks involved a purely visual task that estimated the perceived position of a probe stimulus. To compare a visual probe against a visual reference stimulus, we used a dual adaptation field, in which pointing movements to one region of the visual field were adapted and movements to the opposite field were left unadapted. Any effect of motor adaptation on visual perception should have manifested as a shift in apparent location of the probe stimulus. We found a significant change in visual localization after pointing adaptation only in Experiment 1, in which participants had to slide a stimulus to match the position in depth of a comparison stimulus. After adaptation, participants localized stimuli further in depth, irrespective of the pointing adaptation direction. This result is surprising, as one would expect the visual shift to follow the adaptive shift. For instance, Volcic et al. (2013) induced pointing adaptation with an extended reach feedback. They found that pointing adaptation shifted the position where the object size is estimated accurately towards the adapted location. The direction of their perceptual shift is consistent with our results from extended reach adaptation. However, as we found the same effect in shortened reach adaptation, a simple explanation such as “vision follows pointing adaptation” is not available. We did not find any effect of pointing adaptation on vision in our Experiment 2. In that experiment, we used a task in which two absolute spatial positions had to be compared. This task is different than that of Experiment 1 and that of Volcic et al. (2013) where a comparison position had to be actively matched. Future research could further investigate whether active localization and passive observation are differently affected by sensorimotor adaptation.

An important aspect in this reasoning concerns the status of adaptation. Does it really occur on a motor stage or is it rather a conscious strategy to induce compensation to the artificial feedback error. We strived to make our adaptation procedure to invoke realignment (“true adaptation”) and not the strategic recalibration. First, in order to avoid stereotypic movements, we created two adaptation targets at each side. Second, we introduced the distortion gradually over 66% of the adaptation trials, because previous studies demonstrated that gradual exposure increases the level of adaptation compared to a single-step introduction of the shift (Michel et al., 2007). We also checked whether adaptation affected the pointing planning. We found that peak velocities were increased after shortened reach adaptation and decreased after extended reach adaptation. While this finding does not represent irrevocable evidence for true adaptation, it does at least show that we successfully altered movement plans of participants. In an attempt to reduce the error, participants could have consciously re-planned their pointing movements. However, this hypothesis would predict a uniform generalization of adaptation which stands in stark contrast to our findings.

Our dual field adaptation method was similar to that used in a previous study (Gharamani et al., 1996). They demonstrated a generalization of pointing adaptation that was decaying from the adapted region. In our case this holds true only for the z-direction dimension, whereas in x-direction the aftereffects increase further from the non-adapted field. This difference in generalization might be due to the different adaptation methods in both studies. During adaptation, Gharamani et al. (1996) registered the success of the trial and showed the cursor only if the fingertip was within 0.5 cm of the target. They report that it made pointing difficult and subjects had to move their fingers around and received verbal aid in case it took them long. In our experiment we excluded most of monocular depth cues. Early studies have shown that movement performance drastically decline if they are guided only by monocular cues (Marotta et al., 1997), however others found that movement trajectories might be relatively unaffected by this restriction (Watt & Bradshaw, 2003; Knill & Kersten, 2003). From the remaining binocular cues in our setup, only binocular disparity was a valid indicator of depth, as ocular convergence cannot co-vary with virtual stimulus depth in a head-mounted display. Vergence has measurable supportive effects on depth estimates but binocular disparity clues play the major role (Tresilian et al., 1999).

In summary, our study suggests that pointing movements in depth are biased towards the natural grasping distance. Using pointing adaptation separately in opposite directions, we could dissociate the hypothesis that pointing follows vision from the hypothesis that pointing is biased by a natural grasping distance prior. Adaptation was strongest for those movements that went towards the natural grasping distance, clearly suggesting the latter hypothesis to be true.

